# Genomic and physiological characterization of new Plant Growth Promoting Bacilli isolated from salt-pans

**DOI:** 10.1101/2021.05.17.444429

**Authors:** Claudia Petrillo, Stefany Castaldi, Mariamichela Lanzilli, Matteo Selci, Angelina Cordone, Donato Giovannelli, Rachele Isticato

## Abstract

Massive application of chemical fertilizers and pesticides has been the main strategy used to cope with the rising crop demands in the last decades. The indiscriminate use of chemicals while providing a temporary solution to food demand has led to a decrease in crop productivity and an increase in the environmental impact of modern agriculture. A sustainable alternative to the use of chemicals for crop production is the use of microorganisms naturally capable of enhancing plant growth and protecting crops from pests, known as Plant-Growth-Promoting Bacteria (PGPB). Aim of the present study was to isolate and characterize PGPB from salt-pans sand samples able to ameliorate plant fitness. To survive high salinity, salt-tolerant microbes produce a broad range of compounds with heterogeneous biological activities that are potentially beneficial for plant growth. We have isolated and screened *in vitro* a total of 20 halophilic spore-forming bacteria for phyto-beneficial traits and compared the results with two *Bacilli* recently isolated from the rhizosphere of the same collection site and recently characterized as potential biocontrol agents. Whole-genome analysis on five selected halophilic strains confirmed the presence of numerous gene clusters with PGP and biocontrol functions and of novel secondary-metabolite biosynthetic genes potentially involved in plant growth promotion and protection. The predicted biocontrol potential was confirmed in dual culture assays against several phytopathogenic fungi and bacteria. Interestingly, the absence of predicted gene clusters with known biocontrol functions in some of the isolates was not predictive of the *in vitro* results, supporting the need of combining laboratory assays and genome mining in PGPB identification for future applications.

## 1. Introduction

In the past decades social concern about the environmental effects of the uncontrolled use of chemical pesticides, fertilizers, and herbicides in the agricultural field has risen considerably. The use of chemicals for the protection and enhancement of crops has led to several negative consequences: the formation of stable phytopathogenic variants, the reduction in the number of beneficial microorganisms, and the accumulation of toxic substances in soils and aquatic ecosystems (Reddy et al., 2009; Pertot et al., 2017). Given the increased global demand for crop production, researchers and industries are seeking new, more sustainable and greener approaches to pesticides and fertilizers (Glick et al., 2007). In this framework, the use of microorganisms known as Plant-Growth Promoting Bacteria (PGPB) for crop production and protection appears to be a promising alternative. PGPB improve crop fitness and yields both through direct and indirect mechanisms. Direct mechanisms include the promotion of alternative nutrient uptake pathways, through the solubilization of phosphorus, the fixation of atmospheric nitrogen, the acquisition of iron by siderophores, and the production of growth hormones and molecules, like vitamins, amino acids, and volatile compounds (Babalola, 2010). Indirect mechanisms instead, include the prevention or reduction of the damage induced by phytopathogens through the production of different classes of antimicrobial compounds, such as hydrolytic enzymes that can lyse a portion of the cell walls of many pathogenic fungi (Jadhav et al., 2017).

The work presented here is part of a wider study aimed at identifying and selecting halophilic *Bacilli* with potential applications as biofertilizers or biocontrol agents. For this purpose, samples from the rhizosphere of the nurse plants *Juniperus sabina* and nearby soils were collected from salt-pans (Castaldi et al., 2021). Nurse plants, such as *J. sabina*, exert beneficial effects on their surrounding ecosystem, facilitating the growth and development of other plant species. This positive effect is in part due to the plant influence on the composition of soil microbial communities, generally selecting for microorganisms capable of mineralizing nutrients, enhance soil fertility, and thus to promote plant growth and health (Hortal et al., 2013; Goberna et al., 2014; Rodríguez-Echeverría et al., 2016). For this reason, the nurse-plants rhizosphere and relative surrounding soil are a useful source of PGPB. In addition, bacteria growing in extreme environments, like salt-pans, have developed complex strategies to survive harsh conditions, which include the production of an array of diverse compounds, such as antioxidant pigments, lytic enzymes, and antimicrobial compounds, making them interesting biotechnological targets (Anwar et al., 2020). Among the PGPB, bacteria belonging to the *Bacillus* genus are of particular interest given their resistance to stressful environments and conditions due to their capacity of producing spores (Pesce et al., 2014), together with the ability to release a broad spectrum of secondary metabolites, the easy genetic manipulation, and the great ability to colonize plant surfaces (Kumar et al., 2011). In addition, the effectiveness of halo-tolerant *Bacillus* spp. to increase the growth of various crops under salt stress conditions has been widely reported (Shultana et al., 2020). Recently, we have identified and characterized PGPB *Bacillus* strains isolated from the rhizosphere of *J. sabina* (Castaldi et al., 2021). The two strains, named *Bacillu sp*. RHFS10 and *Bacillu sp*. RHFS18, emerged for their promising PGP traits. These strains produce siderophores and solubilize phosphorus, enhancing plant nutrients uptake, and secrete indoleacetic acid (IAA), a phytohormone playing a key role in both root and shoot development. Additionally, both isolates showed a strong biocontrol activity, inhibiting the fungal phytopathogen *Macrophomina phaseolina* growth (Castaldi et al., 2021).

Here we present the results of the screening of twenty halophilic *Bacilli* isolated from salt-pan sand samples. All the strains were characterized for phyto-beneficial traits and five strains emerged for their high potentiality as biofertilizers and biocontrol agents. Comparative genomic analysis of the five sand strains and the previously characterized rhizospheric strains RHFS10 and RHFS18 were extracted and revealed the presence of known genes involved in plant growth promotion and protection, sustaining, in part, the activities observed *in vitro*.

## 2. Materials and methods

### 2.1 Isolation of bacteria

*Bacillus* strains used in this study were isolated from sand samples collected in the proximity of *J. sabina* plants growing in the salt-pans of Formentera (Spain). Sand samples were heat-treated at 80 °C, to kill vegetative cells and select for spore-forming bacteria, and 1 g of sample was suspended in 9 mL of TY broth (10 g/L tryptone, 5 g/L yeast extract, 8 g/L NaCl) following the heat treatment (Cangiano et al., 2010). Aliquots of supernatant from serial dilutions showing positive growth were spread on TY agar plates and incubated at 30±1 °C for 4-5 days. Pure cultures were obtained by serial sub-culturing and stored at − 80 °C into glycerol stocks (Giglio et al., 2011).

### 2.2 Phenotypic characterization and growth conditions

The phenotypic variants of isolated strains were determined by visual inspection. The facultative anaerobic growth was determined using the AnaeroGen sachets (Unipath Inc., Nepean, Ontario, Canada) placed in a sealed jar with bacteria streaked on TY agar plates and incubated at 37 °C for 3 days. To confirm the sporulation ability, the bacteria were grown in Difco sporulation medium (DSM) (8 g/L Nutrient broth No. 4, 1 g/L KCl, 1 mM MgSO_4_, 1 mM Ca(NO_3_)_2_, 10 μM MnCl_2_, 1 μM FeSO_4_, Sigma-Aldrich, Germany). The optimum growth conditions were determined by growing the isolated strains in TY agar at different pH (2.0, 4.0, 6.0, 7.0, 8.0, 10.0, 12.0) (Cangiano et al., 2014), temperatures (4, 15, 25, 37, 50, 60 °C) (Petrillo et al., 2020) and salinity (0, 5, 10, 13, 15, 18 %) ranges.

### 2.3 Plant Growth-Promoting (PGP) traits

#### 2.3.1 Phosphate solubilization

The phosphate solubilization activity was evaluated by bacteria spot inoculation onto Pikovaskya’s agar medium. The plates were incubated at 28 °C for 10 days. The formation of transparent zones around the bacterial colonies indicates a positive result (Schoebitz et al., 2013).

#### 2.3.2 Siderophores production

The siderophores production was determined by the Chrome Azurol S (CAS) assay. Overnight-grown bacterial cultures were spot-inoculated on CAS agar plates and incubated at 28 °C for 4 days. After 1 hour, the formation of a yellow-orange halo zone around the bacterial colony was a positive indicator of siderophore-production (Pérez-Miranda et al., 2007).

#### 2.3.3 Indoleacetic acid detection

The indoleacetic acid detection (IAA) production was determined with bacteria grown in LB broth at 37 °C for 4 days with shaking at 150 rpm. Following growth, 1 mL of bacteria supernatant was mixed with 2 mL of Salkowski reagent (0.5 M FeCl_3_ in 35 % HClO_4_ solution) incubated at room temperature for 30 min. The formation of pink color indicates IAA production (Damodaran et al., 2013).

#### 2.3.4 Biofilm production and swarming motility

To detect the ability to produce biofilm, 10 µL of fresh bacterial culture were inoculated into 1 mL of sterile LB medium, and the tubes incubated statically at 37 °C for 48 hours (Haney et al., 2018). Swarming motility was tested by spot-inoculating the bacteria strain on LB agar 0.7 % plates and incubated at 37 °C overnight.

### 2.4 Whole-genome sequencing of the selected PGPB

DNA extraction was performed using the DNeasy PowerSoil kit (Qiagen, Hilden, Germany) according to the manufacturer’s instructions. Genome sequencing was performed by MicrobesNG (Birmingham, UK) with the genomic DNA library prepared using the Nextera XT library prep kit (Illumina) following the manufacturer’s protocol. Libraries were sequenced on the Illumina HiSeq using a 250 bp paired-end protocol. Reads were adapted and trimmed using Trimmomatic 0.30 with a sliding window quality cutoff of Q15 (Bolger et al., 2014) and the *de novo* genome assembly was carried out with SPAdes (version 3.7) via MicrobesNG. Genomes were annotated using Prokka (Seemann, 2014). Biosamples accession numbers for strains RHFB, RHF2, RHF6, RHF12, RHF15, RHS10 and RHFS18 are, respectively: SAMN17389615, SAMN17389609, SAMN17389610, SAMN17389612, SAMN17389613, SAMN17389611, SAMN17389614. MIGS compliant details regarding each genome are available in the Supplementary Material Table S1.

Average Nucleotide Identity (ANI) values between the sequenced genomes and the closest bacterial species identified from the 16S rRNA phylogenetic analysis (see below) were obtained using the OrthoANI algorithm of EZBioCloud (Yoon et al., 2017). An ANI similarity of 95 % was considered as a cut-off for species delineation.

### 2.5 Phylogenetic analysis

The 16S rRNA genes were extracted from the sequenced genomes using Anvi’o v2.3.3 (Eren et al., 2021) and compared to 76 reference 16S rRNA genes from closely related strains identified using the Genome Taxonomy Database (GTDB) (https://gtdb.ecogenomic.org) taxonomy and retrieved from the NCBI database. All sequences were aligned using Seaview 4.4.0 software (Corrado et al., 2021) and the phylogenetic tree was constructed using the Maximum-likelihood algorithm with model GTR+I+G4. Statistical support was evaluated by the approximate likelihood-ratio test (aLRT) and is shown at the corresponding nodes of the tree. *Clostridium difficile* is used as an outgroup to root the tree.

### 2.6 Evaluation of potential biocontrol activity

Isolated bacterial strains were tested *in vitro* for growth inhibitory activity against phytopathogenic fungi and bacteria listed in Table 1. Fungi were stored on Potato Dextrose Agar (PDA) in Petri dishes and deposited in the fungal culture collection of the Plant Pathology Department of the University of Buenos Aires (FAUBA, Argentina), except for *S. vesicarium*. Dual-culture plate method was carried out to detect the antifungal activity (Xu and Kim, 2014). Fungal plugs of 6 × 6 mm diameter were placed at the center of PDA plates and 5 µL of bacterial strains grown overnight in TY broth were placed on the opposite four sides of the plates 1.5 cm away from the fungal disc. This method was repeated for each fungus. Controls consisted of plates containing the fungal plugs alone. All plates were incubated at 28 °C for 5-7 days. The antagonism activity against bacterial phytopathogens was performed as described in Li et al. (2020) with some modifications. Bacterial pathogens were streaked on TY plates and incubated at 25 °C overnight. Single colonies were suspended in TY broth and incubated at 25 °C. Approximately 1×10^*− 6*^ CFU/mL were mixed with melted TY agar medium before pouring plates. After solidification, 5 μl of bacterial isolates solution (OD600 = 1.0) was spot-inoculated onto the plates and incubated at 28 °C for 48 h, before measuring the diameters of inhibition halos. All experiments were performed in triplicate.

**Table 1.** List of the phytopathogenic fungi and bacteria used in this study.

| Pathogen type | Species | Strain | Provenience | Host plant |
| --- | --- | --- | --- | --- |
| **Fungi** | *Macrophomina phaseolina* | 201 2013-1 | Argentina | soy |
|  | *Colletotrichum truncatum* | 17-5-5 | Argentina | soy |
|  | *Drechslera teres* | FT | Argentina | barley |
|  | *Cercospora nicotianae* | Ck_2017_B35 | Bolivia | soy |
|  | *Stemphylium vesicarium* |  | Italy | pear |
| **Bacteria** | *Pseudomonas tolaasii* | 2192 | - | mushroom |
|  | *Pseudomonas syringae* pv *tabaci* | ICMP 2706 | - | tobacco |
|  | *Pseudomonas syringae* pv *panici* | ICMP 3955 | - | rice |
|  | *Pseudomonas caryophylli* | NCPPB349 | Italy | carnations |
|  | *Pseudomonas syringae* pv *syringae* | B475 | - | mango |
|  | *Pseudomonas syringae* pv *japonica* | ICMP 6305 | - | wheat |
|  | *Pseudomonas syringae* pv *papulans* | Psp26 | - | apple |

### 2.7 Identification of biosynthesis gene clusters

Obtained genomes were analyzed by antiSMASH 5.0 (Blin et al., 2019) and BAGEL 4 (van Heel et al., 2018) to identify biosynthesis gene clusters (BCGs) of potential antimicrobial compounds such as nonribosomal peptide synthetases (NRPSs), polyketide synthases (PKSs), post-translationally modified peptides (RiPPs), hybrid lipopeptides (NRPS-PKS) and bacteriocins. BGCs that shared less than 70 % amino acid identity against known clusters were regarded as novel.

## 3. Results and discussion

### 3.1 Isolation and characterization of spore-forming Plant Growth Promoting Bacteria (PGPB)

Spore-forming bacteria were specifically isolated from sand samples collected from gaps among nurse plants, belonging to the genus *J. sabina*, in salt-pans as described in the Materials and Methods section. Based on morphological characteristics, a total of 20 isolates were selected and preliminarily characterized for growth properties (Table S2). All the strains can be classified as facultative anaerobic, mesophiles and moderate halophiles, excluding RHF5 strain which survives up to 60 °C and strain RHFB unable to grow at temperature and salt concentration higher than 37 °C and 5 % NaCl, respectively (Ventosa et al., 1998; Schiraldi and De Rosa et al., 2016).

To identify potential PGPB, the 20 strains were evaluated *in vitro* for physiological traits associated with plant growth enhancement and biocontrol ability (Table 2). Strain performance was compared with those of two promising PGPB, RHFS10 and RHFS18 strains, belonging to the *Bacillus* genus and isolated from *J. sabina* rhizosphere of the same collection site (Castaldi et al., 2021) and proposed as biocontrol agents for their antagonistic activity against the phytopathogen *M. phaseolina*. Most of the new strains displayed root-colonization phenotypes since able to surface spread by swarming and to form biofilms (Amaya - Gómez et al., 2020), while only five were found either positive to both solubilization of phosphate, indoleacetic acid (IAA) and siderophore production. Strains RHF6, RHF15, and RHFB showed a better performance than when compared against the already characterized rhizobacteria strains RHFS10 and RHFS18, confirming that the microenvironments created under or nearby nurse shrubs are a promising source of PGPB (Rodriguez-Echeverria et al., 2016). All bacterial isolates were tested for *in vitro* activities of their extracellular hydrolytic enzymes (lipase, protease, amylase, xylanase, cellulase) usually associated with biocontrol activity (Pal and McSpadden Gardener, 2006). As reported in Table 2, the highest hydrolytic activity was observed for RHF12, RHF15, and RHFB strains, comparable with that exerted by rhizosphere strains RHFS10 and RHFS18. Based on these results, five strains sourced from sand samples (RHF2, RHF6, RHF12, RHF15, RHFB) and the two strains from rhizosphere (RHFS10 and RHFS18, Castaldi et al., 2021), showing potential PGP functions, were selected for whole-genome sequencing.

**Table 2.** Summary of plant growth-promoting and biocontrol traits exhibited by 20 spore-forming bacteria isolates.

| Strain | PGPB ACTIVITIES |  |  |  |  | HYDROLYTIC ACTIVITIES |  |  |  |  |
| --- | --- | --- | --- | --- | --- | --- | --- | --- | --- | --- |
|  | Biofilm | Swarming | PVK | IAA | Sideroph. | Lipase | Protease | Amylase | Xylanase | CMC |
| RHF1 | - | ++ | ++ | - | + | - | ++ | ++ | + | ++ |
| **RHF2** | + | + | + | ++ | + | - | + | + | + | + |
| RHF3 | - | - | - | - | - | + | ++ | ++ | + | - |
| RHF4 | - | + | - | - | + | + | ++ | ++ | - | + |
| RHF5 | + | - | - | + | - | - | + | ++ | - | - |
| **RHF6** | + | + | ++ | +++ | ++ | - | + | + | + | ++ |
| RHF7 | + | - | - | - | - | - | + | + | - | - |
| RHF8 | ++ | ++ | - | + | - | - | ++ | ++ | ++ | - |
| RHF9 | - | - | + | + | - | - | ++ | ++ | - | - |
| RHF10 | - | - | - | + | - | - | ++ | + | + | + |
| RHF11 | + | + | - | - | - | - | + | + | + | - |
| **RHF12** | ++ | ++ | + | + | ++ | - | ++ | ++ | ++ | ++ |
| RHF13 | - | ++ | ++ | ++ | + | + | - | ++ | + | ++ |
| RHF14 | - | - | - | - | - | + | + | + | + | - |
| **RHF15** | ++ | ++ | ++ | ++ | ++ | + | ++ | ++ | ++ | ++ |
| RHF16 | - | - | - | - | - | + | + | + | - | - |
| RHF17 | ++ | ++ | + | - | + | + | + | + | ++ | + |
| **RHFB** | + | + | ++ | +++ | ++ | ++ | ++ | ++ | ++ | + |
| RHFE | - | - | - | - | - | + | + | + | - | - |
| RHFL | + | - | - | - | - | - | + | + | - | - |
| **RHFS10<sup>1</sup>** | + | ++ | + | + | ++ | ++ | ++ | ++ | ++ | ++ |
| **RHFS18<sup>1</sup>** | ++ | + | + | + | ++ | + | ++ | ++ | ++ | ++ |

<sup>1</sup>from Castaldi et al., 2021
no activity (-), halo or colony diameter < 5 mm (+), halo or colony diameter ≥ 5 mm (++), halo or colony diameter 10 mm (+++). The strains selected for further studies are indicated in bold. PVK, Pikovskaya; IAA, Indoleacetic acid, CMC, Carboxymethylcellulose.

### 3.2. Genome sequencing and phylogenetic analysis

The obtained genomes had a coverage of ∼30×, with a variable number of contigs between 40 and 1,105 for RHF15 and RHFS18, respectively (Table 3). The genome of strain RHFS18 was particularly fragmented, and repeated sequencing of the same strain did not yield improved assembly suggesting that the results are not dependent on a low quality sequencing library. The obtained genomes are approximately 4 Mbp long except for RHFB’s genome, being the longest (5.6 Mbp) and the one with the highest number of predicted protein coding sequences compared to the others.

**Table 3.** General features of the assembled genomes.

| Analysis Statistics | Strains |  |  |  |  |  |  |
| --- | --- | --- | --- | --- | --- | --- | --- |
|  | RHFB | RHF2 | RHF6 | RHF12 | RHF15 | RHFS10 | RHFS18 |
| Size (bp) | 5,648,75 | 4,003,76 | 4,066,37 | 4,096,20 | 4,232,83 | 4,254,65 | 3,936,40 |
| Number of contigs | 158 | 52 | 156 | 280 | 40 | 46 | 1,105 |
| Mean GC content (%) | 40.57 | 43.74 | 46.3 | 44.01 | 43.39 | 43.95 | 46.14 |
| CDS | 5,413 | 3,988 | 3,901 | 3,997 | 4,282 | 4,182 | 3,87 |
| N50 | 187,761 | 413,215 | 584,325 | 60,229 | 2,184,72 | 1,139,27 | 6,179 |
| N75 | 82,022 | 306,766 | 292,476 | 34,071 | 1,049,73 | 348,257 | 3,118 |
| L50 | 11 | 3 | 2 | 19 | 1 | 2 | 176 |
| L75 | 21 | 6 | 4 | 42 | 2 | 4 | 397 |

Taxonomic identification of the strains was based on the phylogenetic analysis of the 16S rRNA sequence as well as the whole genome Average Nucleotide Identity. All the isolates were identified as members of the genus *Bacillus* (Figure 1) with six strains out of seven clustering into the same clade, and only strain RHFB falling in a different clade. The phylogenetic divergence observed for RHFB from the other strains agrees with the observed differences in physiological traits for this strain (Table S3). Since most *Bacillus* species are phylogenetically close, 16S rRNA analysis is not always exhaustive to obtain an unambiguous assignment (Rooney et al., 2009). To overcome this issue and classify the strains at the species level, whole genome ANI were used (Table 4). Strain RHFB exhibited 96.95 % ANI against the genome of the closest relative *B. frigoritolerans* and was therefore identified as a *B. frigoritolerans* species. Strain RHF2 was identified as *B. subtilis*, based on 99.96 % ANI score. Strains RHF6 and RHFS18 were classified as members of the *B. amyloliquefaciens* species, exhibiting 99.26 % and 98.3 6% ANI, respectively. Strain RHF12 was identified as *B. halotolerans*, based on 98.04 % ANI score, while RHF15 was classified as *B. gibsonii*, showing 99.6 % ANI score. As shown in Table 4, RHFB, RHF12, and RHFS18 strains were univocally matched with the same species, while for RHF2, RHF6 and RHF15 strains the two analyses returned different results. This mismatch between the two methods of classification is due to the poor discrimination between closely related species of the *Bacillus* genus due to their high morphological, biochemical, and genetic similarities (Celandroni et al., 2019). Since taxonomy annotations based on genetic markers, such as the 16S rRNA gene, can give variable results depending on the strain, ANI-based classification has been preferred in this study when showing ANI scores ≥ 95 % (Jain et al., 2018). Based on this, RHF2, RHF6 and RHF15 were identified as *B. subtilis, B. amyloliquefaciens*, and *B. gibsonii*, respectively (Table 4). Only strain RHFS10 could not be classified at the species level due to the low ANI score (93.48 %) when compared with the closest relative, *B. vallismortis* and it was classified as *Bacillus sp*. RHFS10 (Table 3). Further analysis will be required to fill this classification gap.

**Table 4.**
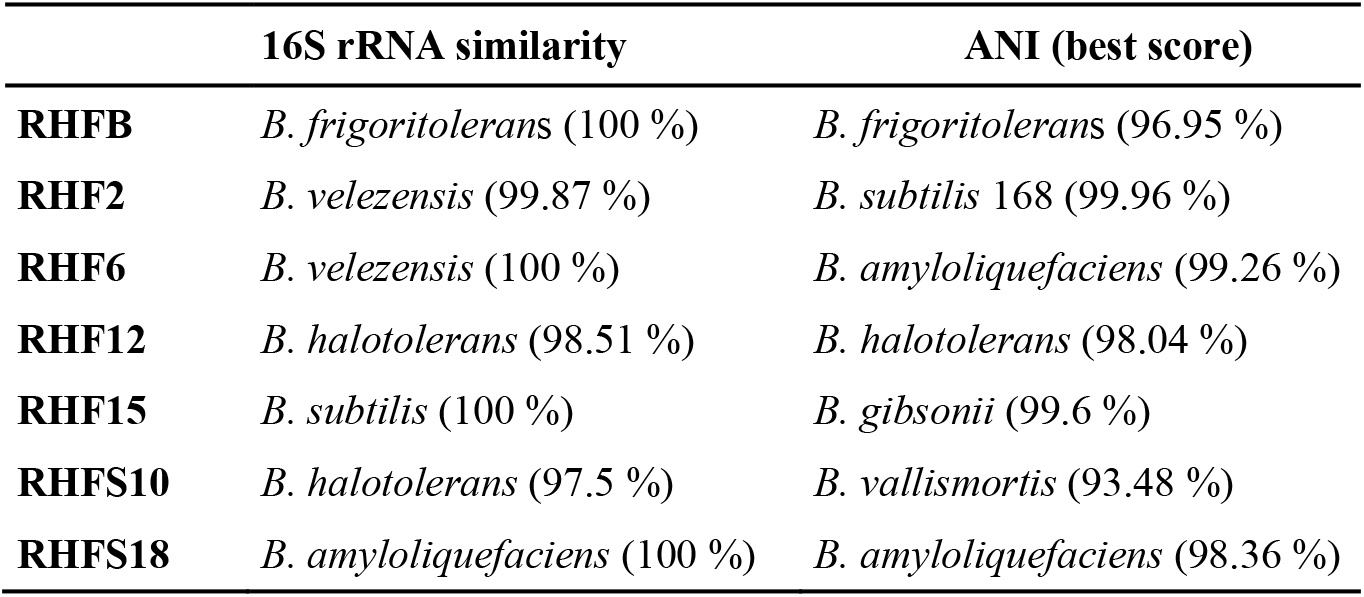
Classification of the seven selected strains. The 16S rRNA similarity and ANI score against the closest relative identified from the phylogenetic analysis are reported for each isolate.

**Figure 1.**
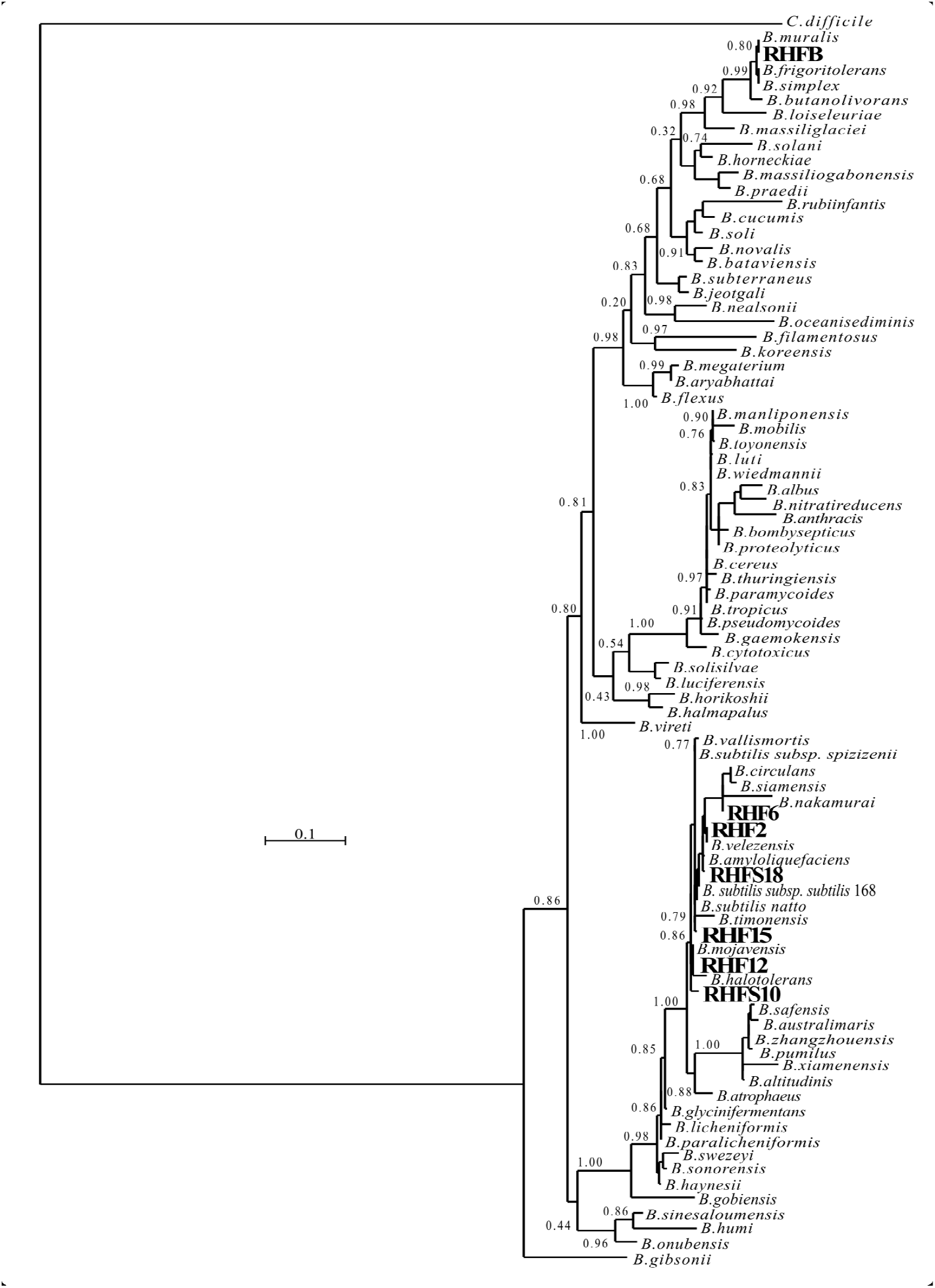
Phylogenetic tree of the spore-forming bacteria isolated from salt-pans. The phylogenetic tree was constructed using the Maximum-likelihood algorithm with model GTR+I+G4, based on 16S rRNA gene sequences. The gene sequences of the isolated bacteria were aligned to reference bacteria belonging to the *Bacillaceae* family according to GTDB. Node support represents the approximate likelihood-ratio test (aLRT) and is shown at the corresponding node of the tree. *Clostridium difficile* is used as an outgroup.

### 3.3. Environmental adaptation to halophilic conditions

The phenotypic plasticity of the salt-pans isolates was investigated by comparing their growth parameters against the closest *Bacillus* species identified by the ANI analysis (Table 4). Temperature, pH and salinity ranges required for growth were evaluated. These parameters are useful to identify distinct phenotypic strategies used by microorganisms to better adapt to environmental conditions (Agrawal, 2001). As expected, taxonomically closer strains showed small differences when compared with each other or with their representative species (red dashed lines in Figure 2). As already highlighted by the phylogenetic analysis, *B. frigoritoleran*s RHFB strain presented a diverging phenotype, especially considering the lower salt tolerance compared to the other isolates. Interestingly, some strains, like *B. halotolerans* RHF12, *B. gibsonii* RHF15, and *B. sp* RHFS10, showed identical growth properties even though belonging to three different *Bacillus* species (Figure 2), while strains of the same species, like *B. amyloliquefaciens* RHF6 and RHFS18, exhibited different adaptations to NaCl concentration and pH range. Moreover, *B. amyloliquefaciens* RHF6 like *B. subtilis* RHF2 were able to grow at higher salt concentrations than their representative species, suggesting an adaptive phenotypic variation to the high salinity condition of salt-pans.

**Figure 2.**
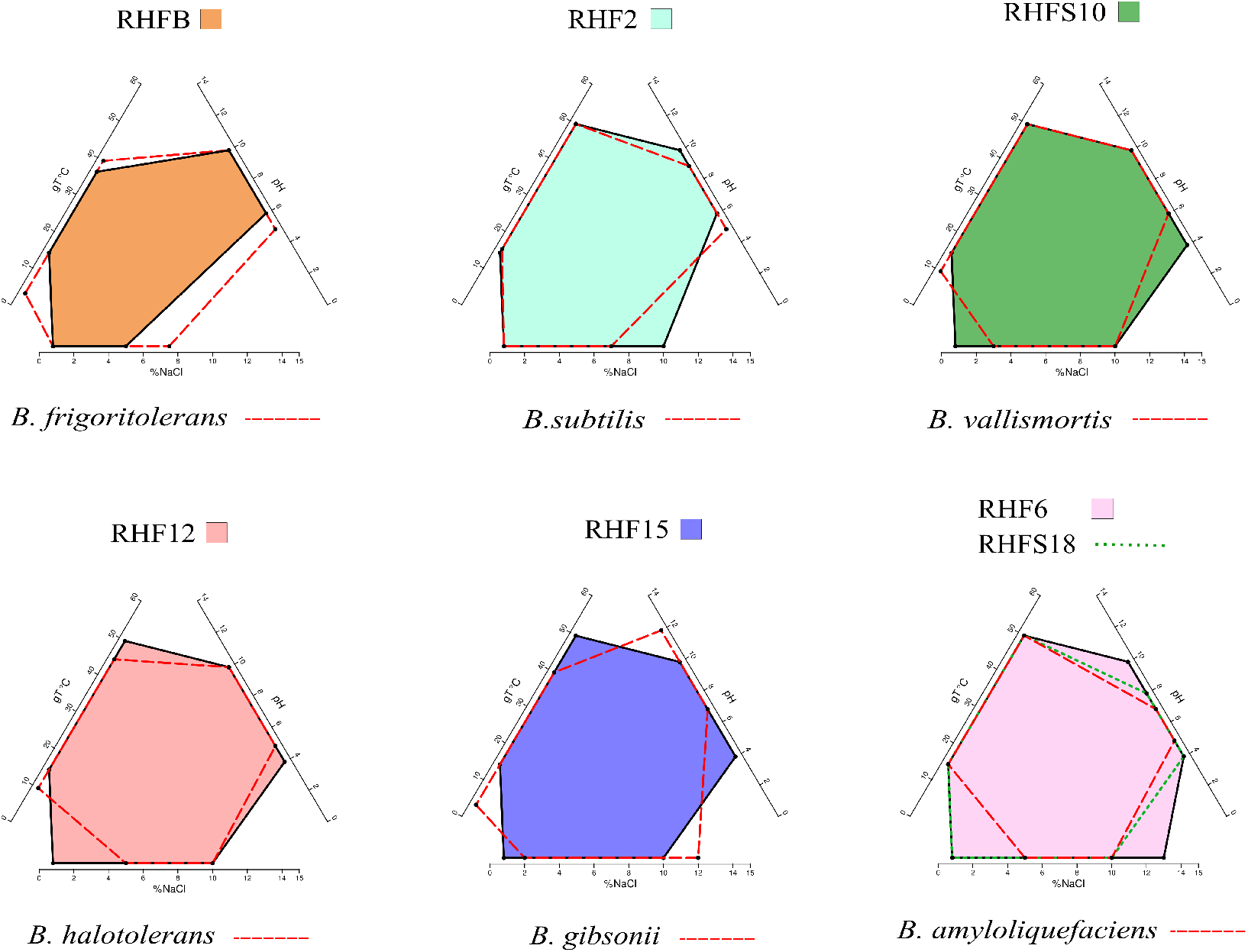
Phenotypic plasticity of the salt-pan isolates. Multivariate polygons plots (Giovannelli et al., in prep) showing the growth temperature (gT °C), pH, and salinity (% NaCl) boundaries observed for the seven isolates (polygons) and the range for the closest relative identified by ANI (red dashed lines). Each edge represents the range for the specific variables projected onto the axis. More information about polygons plot can be found at https://giovannellilab.github.io/polygonsplot/.

### 3.4. Analysis of potential PGP and biocontrol traits

To confirm the *in vitro* PGP characterization of the isolates, a prediction of the genes (Figure 3) and proteins (Table 5) involved in biocontrol activity and plant growth promotion was performed. The analyses identified genes that can be attributed to the strains ability to improve nutrient availability, suppress pathogenic fungi, resist oxidative stress and quorum sensing, in all analyzed genomes. For instance, the genome of all seven strains included the pyrroloquinolone quinone synthase (*pqq*) and the dependent glucose dehydrogenase (*gdh*) genes, involved in mineral phosphate solubilization as well as antifungal activities and systemic resistance induction. Interestingly, both isolates *B. amyloliquefaciens* RHF6 and RHFS18 did not carry the cofactor *pqq* gene cluster, suggesting that other mechanisms could co-exist (Table 2). IAA is one of the most common and effective plant-growth hormones. Besides plants, most rhizobacteria can produce and secrete IAA, increasing the growth and the yield of crops (Bunsangiam et al., 2019). All the strains produced Tryptophan-2-monooxygenase and Indole-3-acetamide hydrolase, able to convert Tryptophan in Indole-3-acetamide and then in IAA, respectively (Bunsangiam et al., 2019). The presence of other tryptophan synthases orthologs (subunits a and b) in all the analyzed genomes suggests alternative IAA biosynthesis pathways potentially involving different intermediates.

**Figure 3.**
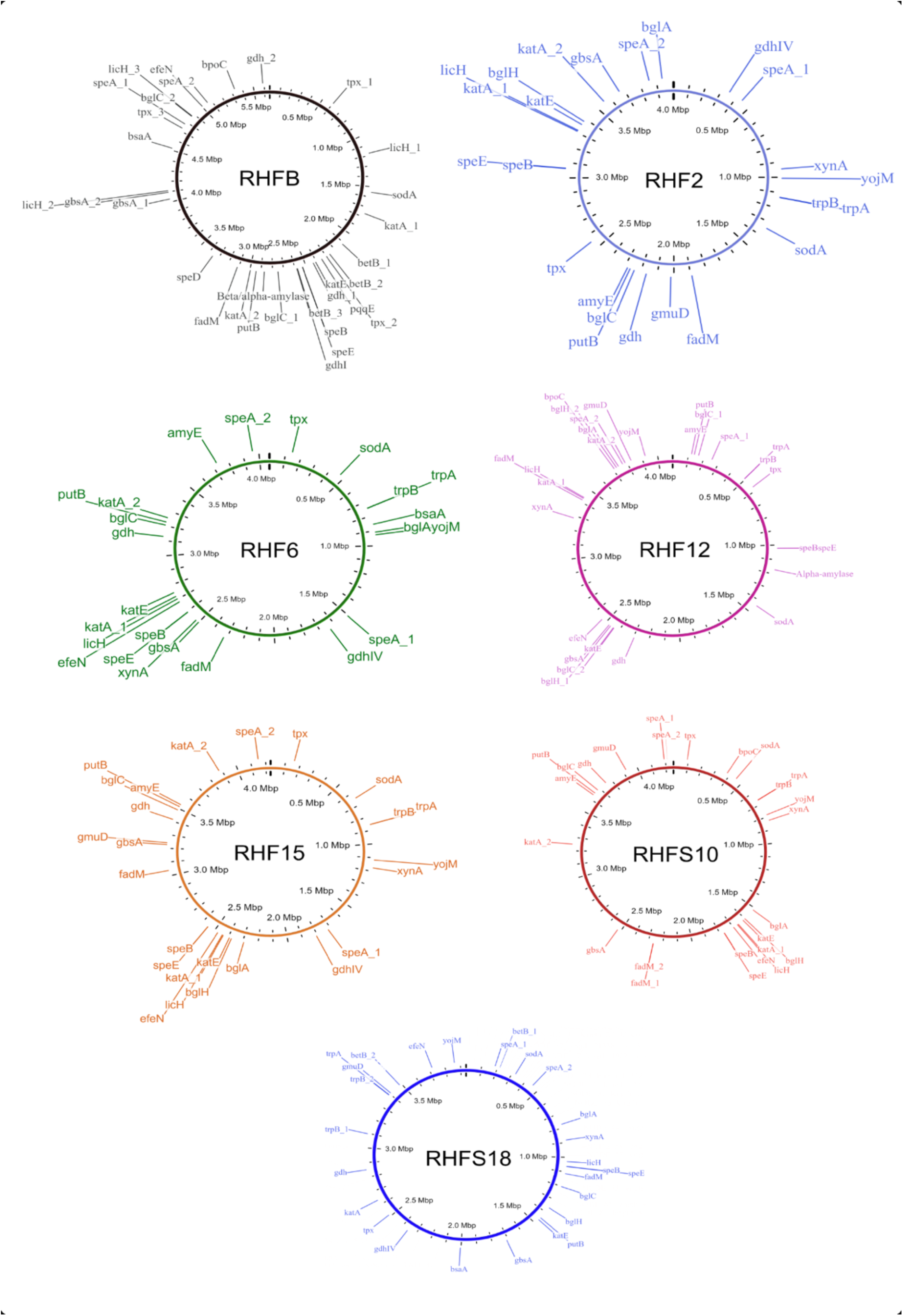
Whole genome representations of the seven isolates showing the location of the identified PGP trait genes.

**Figure 4.**
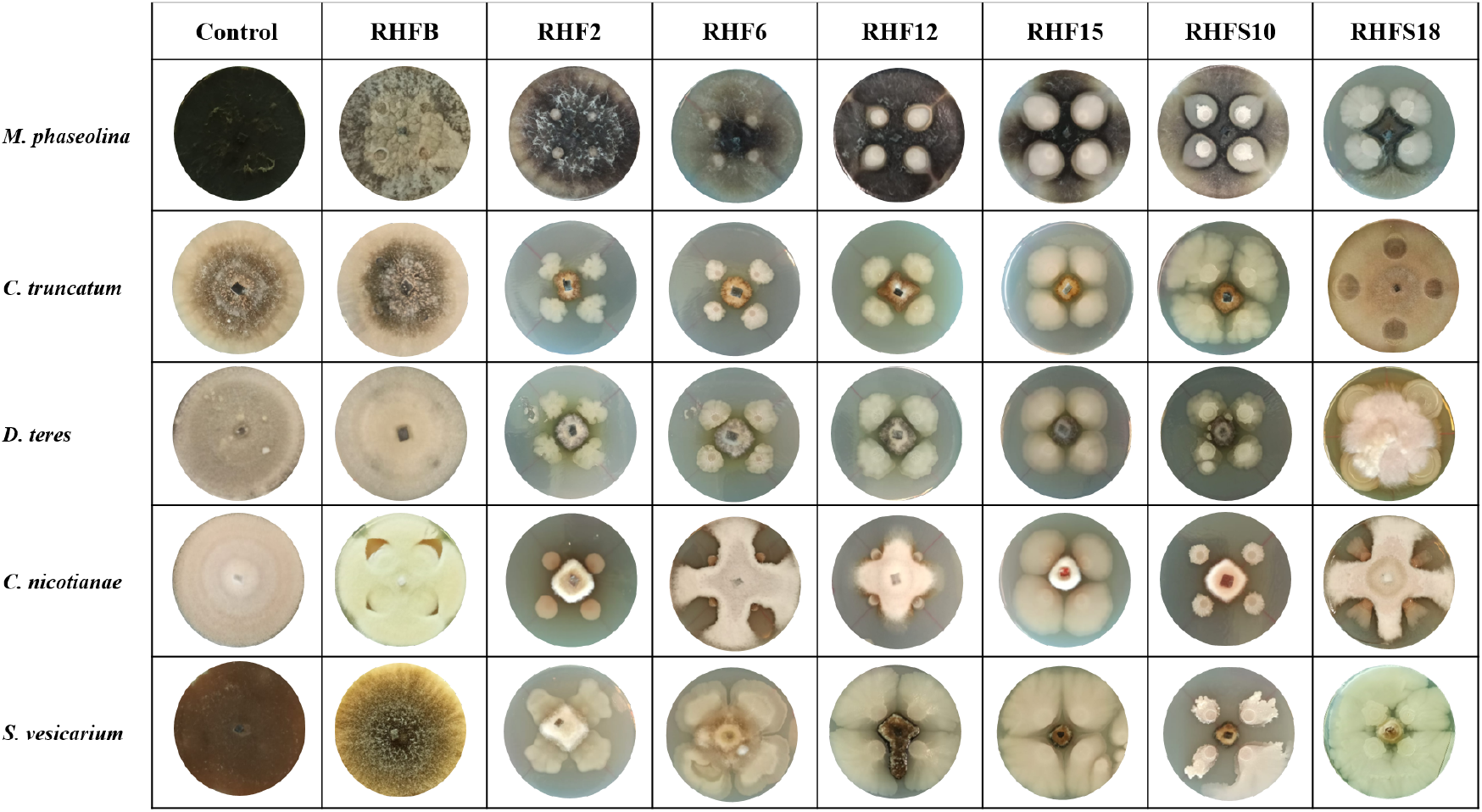
Representative photographs of dual culture assay for *in vitro* mycelial growth inhibition of fungal phytopathogens.

**Table 5.** Plant-Growth-Promoting traits-associated proteins identified in the proteome of the selected strains and their abundance.

| PGP Trait | Gene | RHFB | RHF2 | RHF6 | RHF12 | RHF15 | RHFS10 | RHFS18 |
| --- | --- | --- | --- | --- | --- | --- | --- | --- |
| **Phosphate solubilization** | Pyrroloquinoline quinone (*pqq*) | 1 | 1 | 0 | 1 | 1 | 1 | 0 |
|  | Glucose 1-dehydrogenase (*gdh*) | 2 | 2 | 2 | 2 | 2 | 2 | 2 |
| **Nitrogen fixing** | Nitrogenases | 6 | 6 | 4 | 6 | 6 | 6 | 2 |
| **Nitric oxide synthesis** | Copper-containing nitrite reductase | 1 | 2 | 1 | 3 | 2 | 2 | 1 |
| **IAA biosynthesis** | Indole-3-pyruvate decarboxylase | 1 | 0 | 0 | 0 | 0 | 0 | 0 |
|  | Tryptophan 2-monooxygenase | 4 | 2 | 1 | 2 | 2 | 3 | 2 |
|  | Tryptophan synthase<br>(subunit a and b) | 6 | 7 | 6 | 7 | 7 | 5 | 7 |
|  | Tryptophan aminotransferase | 0 | 0 | 0 | 0 | 0 | 0 | 0 |
|  | Tryptophan decarboxylase | 0 | 0 | 0 | 0 | 0 | 0 | 0 |
|  | Indole-3-acetamide hydrolase | 0 | 0 | 0 | 0 | 0 | 0 | 0 |
| **Putrescine and Spermidine**<br>**-related production** | Arginine decarboxylase | 3 | 2 | 2 | 2 | 2 | 2 | 2 |
|  | Agmatine ureohydrolase | 1 | 1 | 1 | 1 | 1 | 1 | 2 |
|  | Ornithine decarboxylase | 0 | 0 | 0 | 0 | 0 | 0 | 0 |
|  | SAM decarboxylase | 1 | 1 | 1 | 1 | 1 | 2 | 1 |
|  | Spermidine synthase | 1 | 1 | 1 | 1 | 1 | 1 | 1 |
| **ACC deaminase activity** | ACC deaminase | 2 | 2 | 3 | 1 | 2 | 1 | 3 |
|  | D-cysteine desulphydrase | 1 | 0 | 1 | 0 | 0 | 0 | 1 |
| **Antioxidant activity** | Peroxidases | 9 | 10 | 4 | 9 | 9 | 8 | 4 |
|  | Catalases | 10 | 12 | 11 | 11 | 12 | 11 | 8 |
|  | Superoxide dismutase | 7 | 5 | 5 | 6 | 5 | 5 | 5 |
|  | Glutathione peroxidase | 1 | 1 | 1 | 1 | 2 | 1 | 1 |
|  | Glutathione reductase (*gr*) | 0 | 0 | 0 | 0 | 0 | 0 | 0 |
|  | Glutathione S-transferase | 2 | 5 | 2 | 2 | 5 | 2 | 3 |
|  | Choline dehydrogenase | 0 | 1 | 1 | 2 | 2 | 1 | 1 |
|  | Betaine-aldehyde dehydrogenase | 5 | 2 | 2 | 2 | 2 | 2 | 2 |
|  | Proline dehydrogenase | 2 | 3 | 2 | 2 | 2 | 2 | 2 |
| **Cell wall and degrading** | β-glucosidase | 1 | 3 | 2 | 5 | 3 | 3 | 3 |
| | α-glucosidase | 3 | 4 | 4 | 3 | 4 | 4 | 2 |
| | Endo-1,4-β-xylanase | 5 | 7 | 7 | 4 | 5 | 7 | 9 |
|  | Glucosylase | 0 | 0 | 2 | 0 | 0 | 0 | 1 |
| | α-amylase | 0 | 1 | 1 | 1 | 1 | 1 | 1 |
|  | Chitinase | 1 | 0 | 0 | 0 | 0 | 1 | 1 |
| | β-1,3-glucanase | 2 | 3 | 2 | 1 | 2 | 2 | 1 |
|  | Cellulase | 0 | 3 | 2 | 3 | 3 | 4 | 2 |
|  | Protease | 4 | 3 | 3 | 3 | 3 | 2 | 1 |

Only $\geq 40$ % similarity scores were considered. IAA, Indole-3-acetic acid; ACC, 1-aminocyclopropane-1-carboxylate

**Table 6.** Antimicrobial activity of the seven selected strains against phytopathogenic fungi and bacteria.

| Pathogen types | Species | RHFB | RHF2 | RHF6 | RHF12 | RHF15 | RHFS10 | RHFS18 |
| --- | --- | --- | --- | --- | --- | --- | --- | --- |
| **Fungi** | *M. phaseolina* | - | - | - | + | ++ | +++ | +++ |
|  | *C. truncatum* | - | - | +++ | +++ | +++ | +++ | +++ |
|  | *D. eres* | - | - | +++ | +++ | +++ | +++ | +++ |
|  | *C. nicotianae* | - | +++ | ++ | ++ | +++ | +++ | ++ |
|  | *S. vesicarium* | - | ++ | +++ | ++ | +++ | +++ | - |
| **Bacteria** | *P. tolaasii* | - | - | + | - | - | + | - |
|  | *P. syringae* pv *tabaci* | - | ++ | ++ | - | - | + | - |
|  | *P. syringae* pv *panici* | - | ++ | ++ | - | - | + | - |
|  | *P. cariphilly* | - | - | - | - | + | + | - |
|  | *P. syringae* pv *syringae* | - | + | + | - | - | ++ | - |
|  | *P. syringae* pv *japonica* | - | ++ | ++ | - | - | + | - |
|  | *P. syringae* pv *papulans* | - | - | - | - | - | - | ++ |

no inhibition (-), inhibitory zone < 5 mm (+), inhibitory zone 5 mm (++), inhibitory zone >5 mm (+++).

This hypothesis is supported by the observation that *B. frigoritoleran*s RHFB, one of the best IAA producers among the isolated PGPB,

### 3.5. Analysis of potential PGP and biocontrol traits

To confirm the *in vitro* PGP characterization of the isolates, a prediction of the genes (Figure 3) and proteins (Table 5) involved in biocontrol activity and plant growth promotion was performed. The analyses identified genes that can be attributed to the strains ability to improve nutrient availability, suppress pathogenic fungi, resist oxidative stress and quorum sensing, in all analyzed genomes. For instance, the genome of all seven strains included the pyrroloquinolone quinone synthase (*pqq*) and the dependent glucose dehydrogenase (*gdh*) genes, involved in mineral phosphate solubilization as well as antifungal activities and systemic resistance induction. Interestingly, both isolates *B. amyloliquefaciens* RHF6 and RHFS18 did not carry the cofactor *pqq* gene cluster, suggesting that other mechanisms could co-exist (Table 2). IAA is one of the most common and effective plant-growth hormones. Besides plants, most rhizobacteria can produce and secrete IAA, increasing the growth and the yield of crops (Bunsangiam et al., 2019). All the strains produced Tryptophan-2-monooxygenase and Indole-3-acetamide hydrolase, able to convert Tryptophan in Indole-3-acetamide and then in IAA, respectively (Bunsangiam et al., 2019). The presence of other tryptophan synthases orthologs (subunits a and b) in all the analyzed genomes suggests alternative IAA biosynthesis pathways potentially involving different intermediates. This hypothesis is supported by the observation that *B. frigoritoleran*s RHFB, one of the best IAA producers among the isolated PGPB, possessed the indole-3-pyruvate decarboxylase gene, a key enzyme of another Trp-dependent pathway for IAA production.

Finally, all the strains were predicted to be potentially able to fix nitrogen and produce nitric oxide, both useful features in agricultural practices (Ahmad et al., 2013), and to synthesize polyamines, as spermidine and putrescine, and the ACC deaminase, involved in lateral root development and plant growth enhancement under abiotic stress (Xie et al., 2014; Gupta and Pandey, 2019).

As expected, the genome of all the isolated halophilic *Bacillus* strains contained multiple genes involved in antioxidant response, such as peroxidases, catalases, superoxide dismutase and glutathione peroxidase (Hassan et al., 2020) (Table 5). Other enzymes involved in abiotic stress responses were identified in the strains, as the osmoprotectants choline dehydrogenase, betaine-aldehyde dehydrogenase and proline dehydrogenase (Table 5). The predicted production of osmotically active metabolites, as well as ROS scavenging enzymes, reflects the ability of the selected strains to survive in extreme environments, as salt-pans.

Finally, all the isolates possessed in their genomes genes encoding for hydrolases involved in fungal cell-wall and starch degrading pathways, confirming the results obtained with the *in vitro* analysis, except for strain *B. frigoritoleran*s RHFB whose genome did not carry α-amylase or cellulase genes.

### 3.6. Antimicrobial activity screening

To verify the antagonistic potential that emerged from the genome-mining, the isolates were dually cultured with fungal and bacterial plant pathogens (see Table 1 for a list of the used phytopathogens). The results reveal that isolates inhibited plant pathogens growth on plates with different efficiency. Strains *B. subtilis* RHF2, *B. amyloliquefaciens* RHF6, and *B. sp* RHFS10 showed a broad inhibitory spectrum, being able to antagonize both phytopathogenic fungi and bacteria, while *B. halotolerans* RHF12 and *B. amyloliquefaciens* RHFS18 exhibited an antimicrobial activity limited to fungi. The highest antagonistic activity was observed for strain *B. sp* RHFS10, capable of inhibiting the growth of most of the test pathogens, confirming its biocontrol potential already observed by Castaldi et al. (2021). Unexpectedly, *B. frigoritolerans* RHFB exhibited no activity at all. Nevertheless, in the last decade, this species has been identified as a potential insect pathogenic bacterial species, with nematicidal activity (Selvakumar et al., 2011). The diversity observed in the antimicrobial activity against plant pathogens highlighted the phenotypic diversity of sand and rhizosphere isolated *Bacilli*, suggesting that in nature plant-associated bacteria may encounter different phytopathogens that may induce the acquisition of different antagonistic activity.

### 3.7. Genome mining for Bioactive Gene Clusters

The biocontrol potential and the ability to enhance plant growth of PGPB are mostly attributed to their bioactive secondary metabolites. Proteins and metabolites released in the soil by PGPB, indeed, are implicated in root colonization, as well as in interactions with the plant immune response and the surrounding niche (Lugtenberg and Kamilova, 2009; Pieterse et al., 2014; Jamali et al., 2020). The strong antimicrobial activity of selected *Bacillus* strains is most likely due in part to the production of hydrolytic enzymes and siderophores observed *in vitro* assays and confirmed by genome analysis (Table 2 and 5). To better investigate this antagonistic activity, the biosynthetic potential of the halophilic PGPB was evaluated by using antiSMASH 6.0.0 to predict both characterized and unknown functioned secondary metabolites (Figure 5).

**Figure 5.**
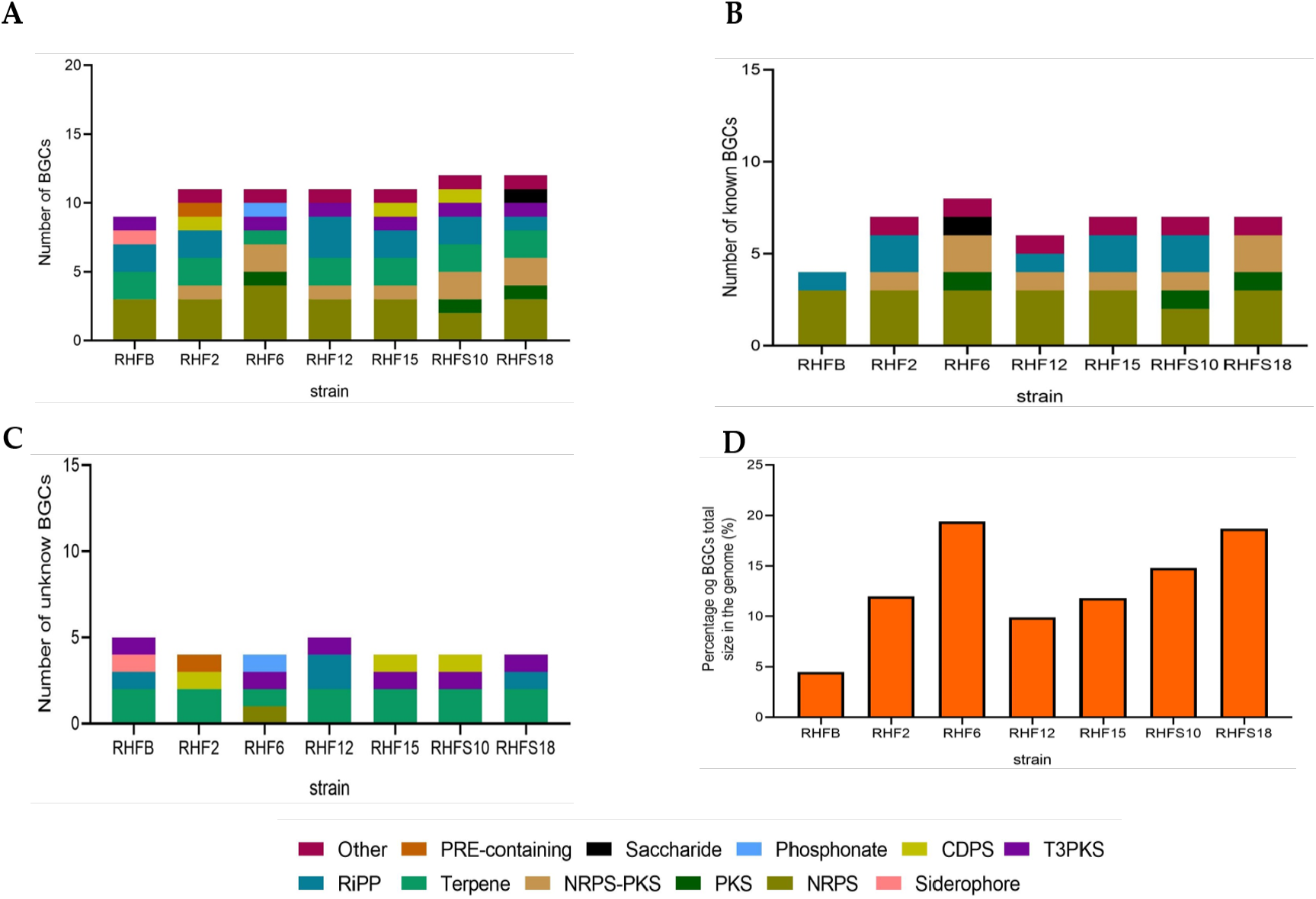
Number of biosynthetic gene clusters harbored by the strains and the percentage contribution of BGCs to the total genome size. (A) Total number of BGCs; (B) number of reported BGCs in the genomes; (C) number of unknown BGCs. BGCs that have different numbers of genes or show less than 70 % protein identity to the reported ones were regarded as novel; (D) the percentage contribution of BGCs to the genomes.

The bacterial isolates harbored Bioactive Gene Clusters (BGCs) coding for nonribosomal peptide synthetases (NRPSs), polyketide synthases (PKSs), post-translationally modified peptides (RiPPs), hybrid lipopeptides (NRPS-PKS) (Figure 5A) and the majority of the BGCs are assigned to known products (Figure 5B, Table S4). The unknown BGCs are type 3 polyketide synthase (T3PKS), RiPPs and terpenes (Figure. 5C, Table S4). The genome of *B. frigoritolerans* RHFB is approximately 5.6 Mbp, and the BGCs account for 4.5 % of the total size (Figure 5D). This strain devotes the lowest percentage of its genome to the synthesis of BGCs with respect to the other selected strains, and the genetic analysis revealed a high abundance of unknown compounds such as RiPPs, T3PKS, and Siderophores (Figure 5C). *B. subtilis* RHF2 and *B. gibsonii* RHF15 devote around 12 and 11.8 % of their genomes to synthesize antimicrobial metabolites, respectively (Figure 5D). These strains synthesize equal numbers of BGCs (13 %) and many of them are known as NRPS, NRPS-PKS and RiPPs (Figure 5B). 4 % of BGCs (Terpene, RiPP and cyclodipeptide synthase CDPS) of both strains are unknown (Figure 5C). This result is similar to the estimation of *B. halotolerans* RHF12, which is 9.9 % of its genome to synthesize BGCs. Both *B. amyloliquefaciens* RHF6 and RHFS18 use 19.4 % and 18.7 % of their genomes, respectively. These bacteria synthesize the highest number of BGCs with respect to the other strains (Figure 3D). 8 % and 7 % of BGCs in strains RHF6 and RHFS18 respectively, are known (Figure 5B) and 4 % in both strains are unknown. Most of the unknown BGCs from the RHF6 strain are NRPS, Terpene, T3PKS, and Phosphonate. In the RHFS18 strain, the abundance of unknown BGCs are represented by Terpene, RiPPs, and T3PKS (Figure 5C). Finally, *B. sp* RHFS10 devotes 14.8 % of its genome to synthesize BGCs (Figure 5D), of which 7 % are known and the most abundance is represented by NRPS and RiPPs (Figure 5B). Interestingly, this strain has the same number and type of unknown BGCs as strain RHF15 represented by Terpene, T3PKS and CDPS (Figure 5C).

### 3.8. Novel Nonribosomal Peptide Synthetases and bacteriocins

NRPs are modular enzymes that synthesize secondary metabolites, some of which are known to be involved in plant disease control (Ongena and Jacques, 2008). Several bioactive compounds produced by *Bacillus* strains fit in this category, such as surfactin or fengycin (Keswani et al., 2020), both of them exhibiting antimicrobial activity potentially exploited for biocontrol in agriculture. We have identified one novel BGC belonging to the class of the NRPs from *B. amyloliquefaciens* RHF6 (Figure 6). This cluster of 66.3 Kb has 6 genes encoding 25 domains, which include 6 condensation (C) domains, 7 adenylation (A) domains, 1 coenzyme A ligase (CAL) domain, 2 epimerization (E) domains, 1 thioesterase (TE) domain, 1 heterocyclization (Cy) domain and 7 peptidyl carrier protein (PCP) domains. Among them, 24 domains are essential components of this cluster, and catalyze the incorporation of 7 amino acids into the final product exhibiting the following sequence: D-Cys–Ser– Cys–Ala–Asn–D-Asn. This cluster shows no similarity to any known BGCs reported in the antiSMASH database (Table S4). The single heterocyclization (C) domain in the first module of the BGC, could form a thiazoline ring from a residue of cystine (Cys). Interestingly, many antimicrobial drugs expose a thiazoline ring (Desai et al., 2016). This allows us to speculate on the potential antimicrobial activity of the compound produced by this novel BGC. The seven genomes were also mined for potential novel bacteriocins BGCs using BAGEL4. Bacteriocins are ribosomally synthesized antimicrobial peptides, generally active against bacteria closely related to producers (Cotter et al., 2013), and classified into three main classes: class I comprehends ribosomally produced and post-translationally modified peptides (RiPPs); class II unmodified peptides, and class III large antimicrobial peptides (Zhao and Kuipers, 2016).

**Figure 6.**
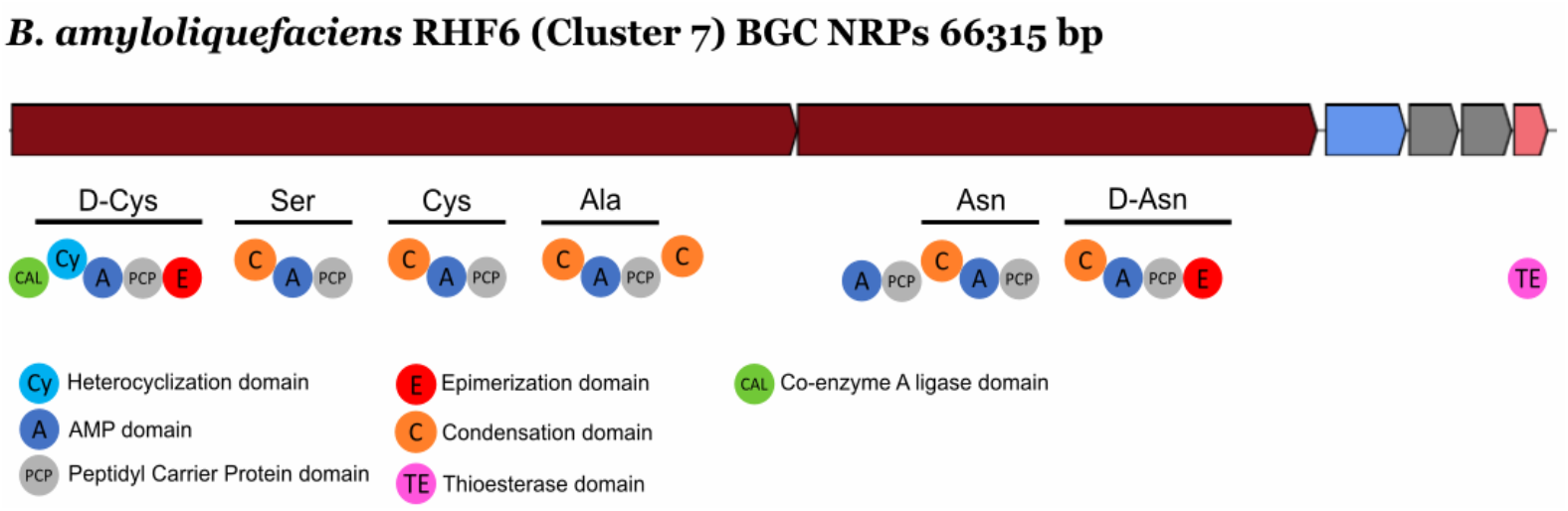
Novel NRP identified from the isolate *B. amyloliquefaciens* RHF6.

These molecules are directed against competitive microorganisms, and therefore generate a selective advantage for the producers. Generally, bacteriocins are highly specific against their target, although some might have a wider spectrum (Jack et al., 1995). The analysis made using BAGEL4, returned 15 regions of interest (in contrast with the antiSMASH analysis which revealed a higher number of bacteriocins Table S4), even though only 6 of them could be classified as novel bacteriocins, sharing ≤ 70 % of similarity with known sequences from BAGEL4 database (Figure 7).

**Figure 7.**
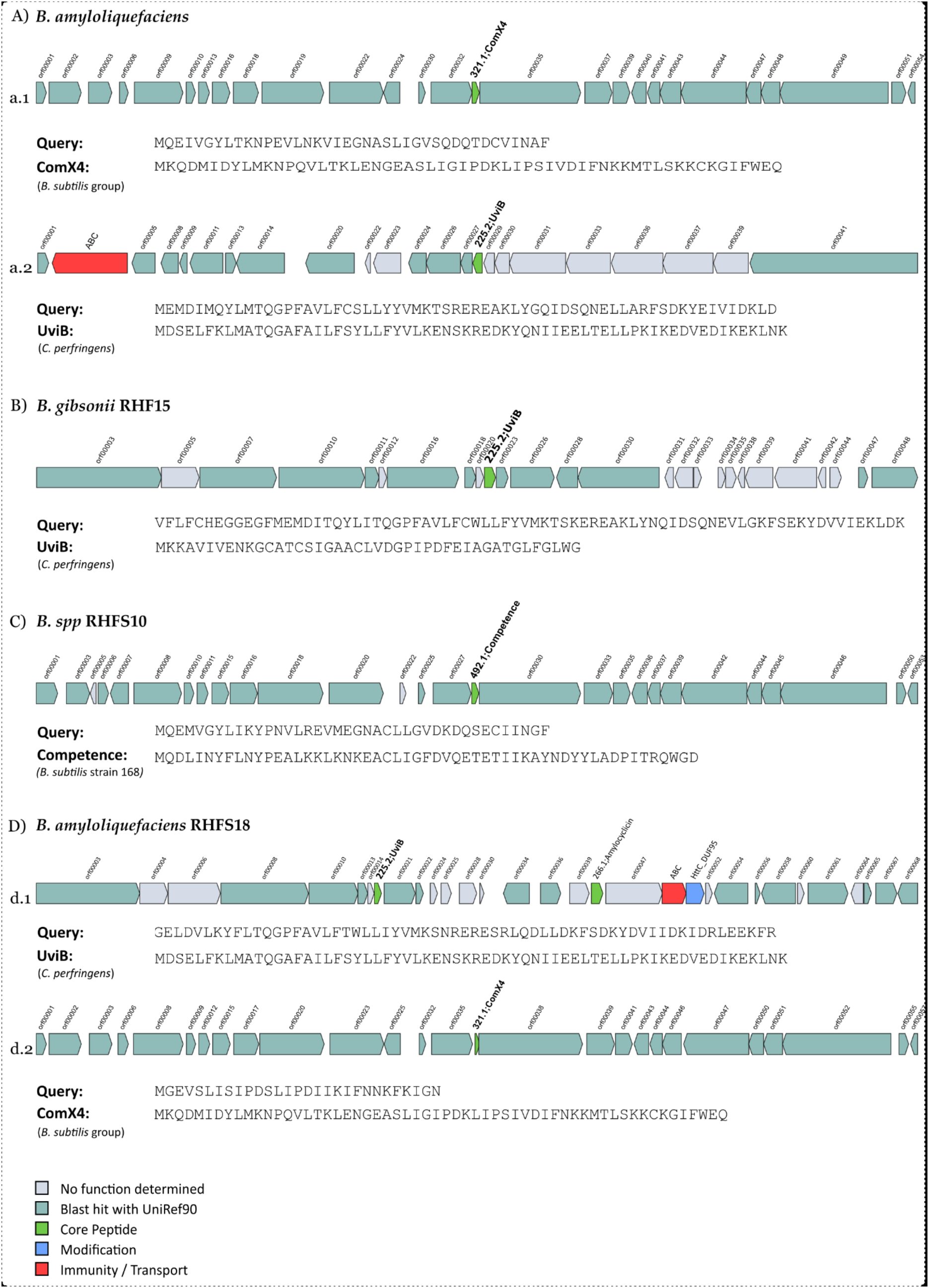
Novel bacteriocins identified from the isolated *Bacillus* strains. The BGCs identified from BAGEL4 analysis are shown and compared to the most similar available in BAGEL4 database.

One orphan BGC of 27 genes is carried by both *B. amyloliquefaciens* RHF6 and RHFS18 strains (Figure 7a.1, 7d.1), although the core biosynthetic genes encode two different precursor peptides of 40 and 29 amino acids, respectively, sharing 41.03 % and 57.14 % of similarity with ComX4 from the *B. subtilis* group. In particular, ComX4 belongs to the ComX subclass of RiPPs according to the BAGEL4 database and it is part of a major quorum-sensing system that regulates the development of genetic competence (Okada et al., 2005) and the production of surfactins (Caulier et al., 2019). *B. amyloliquefaciens* RHF6 also harbors a BGC of 23 genes (Figure 7a.2), with the core biosynthetic gene encoding a 63-amino acids precursor peptide, showing a similarity of 36.51 % compared to UviB, a class II bacteriocin first identified in the mobilizable plasmid pIP404, from *C. perfringens*, known to be bacteriocinogenic (Garnier and Cole, 1988). Interestingly, two different BGCs containing the same gene encoding for a putative UviB-like bacteriocin, were found in strains *B. gibsonii* RHF15 (Figure 7B) and *B. amyloliquefaciens* RHFS18 (Figure 7d.1). Their precursor peptides share 42.1 % and 33.4 % similarity with UviB.

Finally, *B. sp* RHFS10 carries a orphan 28 genes BGC with a core biosynthetic gene encoding a 40-amino acids peptide sharing 35 % of similarity with the competence pheromone of *B. subtilis 168*, a RiPP belonging to class I bacteriocins. *Bacillus* species are known to synthesize many well-studied bacteriocins, such as subtilin, ericin, paenibacillin, subtilosin, thuricin and coagulin (Abriouel et al., 2011). Anyway, it is impossible to predict if the six compounds produced by strains *B. amyloliquefaciens* RHF6, and RHFS18, *B. gibsonii* RHF15 and *B. sp* RHFS10 have antimicrobial properties from genome sequence data only. Despite this, the antagonistic assays *in vitro* suggest that some of them might have antibacterial and/or antifungal activities. This will need to be validated by further experiments.

## 4. Conclusions

In a historic moment in which the increasing population coupled with land degradation aggravates crop production, the use of plant growth promoting bacteria to ensure agricultural productivity has a huge impact on our society. These soil microorganisms enhance plant performance and represent an eco-friendly alternative to chemical fertilizers and pesticides (Hashem et al., 2019). When applied directly to the soil, PGPB promote plant growth by different action mechanisms such as the production of different phytohormones, accelerating the mineralization of organic matter and improving the bioavailability of the nutrients, and protecting plants from pests damages. The beneficial activity exerted by PGPB is in part mediated by a broad spectrum of secondary metabolites and enzymes. For example, polyamines, such as spermidine, play important physiological and protective roles in plants, resulting in an increase in biomass, altered root architecture, and elevated photosynthetic capacity. Until recently, these key metabolites were uncovered only by systematic investigation or by serendipity, often understating the PGPB potentiality during their screening. Many genes involved in PGB activity, in fact, could be silent under standard laboratory conditions, due to the absence of appropriate natural triggers or stress signals. More recently, the onset of the genomic era has facilitated the discovery of these ecologically important metabolites and novel strategies became available for PGPR characterization.

For example, genome mining allows to look over the whole genome of a PGPB strain and highlights genes encoding beneficial enzymes, involved in the enhancement of plant nutritional uptake or modulation of hormone levels, as well as for antimicrobial-encoding BGCs.

In this work, we have isolated soil halophilic *Bacilli* and performed their screening for PGP traits by using standard laboratory procedures and whole-genome analysis. *Bacilli* represent a significant fraction of the soil microbial community and some species are categorized as PGPB (Cazorla et al., 2007). They are also able to produce endospores, which besides enduring harsh environmental conditions fatal for other cell forms (Petrillo et al., 2020), permit easy formulation and storage of commercial PGPB-based products. In addition, salt-tolerant PGPB can easily withstand several abiotic stresses and ameliorate plant growth in degraded soil.

Seven *Bacillus* strains have been selected for *in vitro* PGP traits and identified at the species level by genome analysis. Based on genome mining, not only have we confirmed the beneficial activities PGP found by in vitro analysis, identifying the involved genes, but we have highlighted their strong potentiality by the discovery of novel biosynthesis gene clusters. Our results demonstrated that the genomic analyses, as genome mining, allow a full investigation of PGPB biosynthetic capacity for secondary metabolites and proteins and represent useful tools in the characterization of plant beneficial bacteria. Nevertheless, the divergences observed between the predicted biocontrol functions by found gene clusters and the results obtained by *in vitro* analysis, highlight the need of combining laboratory-assays and genome-mining in identification of new PGPB for future applications.

## Acknowledgments

We thank Marcelo Anibal Carmona (Facultad de Agronomía, Cátedra de Fitopatología, Universidad de Buenos Aires, Buenos Aires, Argentina) to supply the phytopathogenic fungi (M. phaseolina, C. truncatum; C.nicotianae; D.teres) used in this study.

## Data Availability Statement

The genomes generated for this study have been deposited in NCBI under Biosample accession numbers SAMN17389615, SAMN17389609, SAMN17389610, SAMN17389612, SAMN17389613, SAMN17389611, SAMN17389614 for strains RHFB, RHF2, RHF6, RHF12, RHF15, RHS10 and RHFS18, respectively.

## Author Contributions

Conceptualization, R.I.; methodology, S.C. and C.P.; validation, and formal analysis, S.C., C.P., M.L., and M.S.; investigation, S.C., C.P. and D.G.; data curation, S.C., C.P., M.S., A.C. and R.I.; writing original draft preparation, R.I., S.C., C.P. and D.G.; supervision, R.I.; project administration, R.I.; funding acquisition, R.I. All authors have read and agreed to the published version of the manuscript.

## Supplemental material

**Table S1.** The minimum information about genome sequences (MIGS).

| MIGS ID | Property | Term |
| --- | --- | --- |
| MIGS-31 | Finishing quality | High-Quality Draft |
| MIGS-28 | Libraries used | Illumina MiSeq Reagent Kit v2 2x250bp paired-end reads |
| MIGS-29 | Sequencing platforms | Illumina MiSeq |
| MIGS-31.2 | Fold coverage | 30x |
| MIGS-30 | Assemblers | SPAdes version 3.7 |
| MIGS-32 | Gene calling method | Microbial Genome Annotation Pipeline (MiGAP) |
|  | Genbank ID | SAMN17389615 (RHFB), SAMN17389609 (RHF2), SAMN17389610 (RHF6), SAMN17389612(RHF12), SAMN17389613 (RHF15), SAMN17389611 (RHFS10), SAMN17389614 (RHFS18) |
|  | Genbank Date of Release | 03/09/2021 |
|  | BIOPROJECT | PRJNA693507 |
|  | Project relevance | Industrial |

**Table S2.**
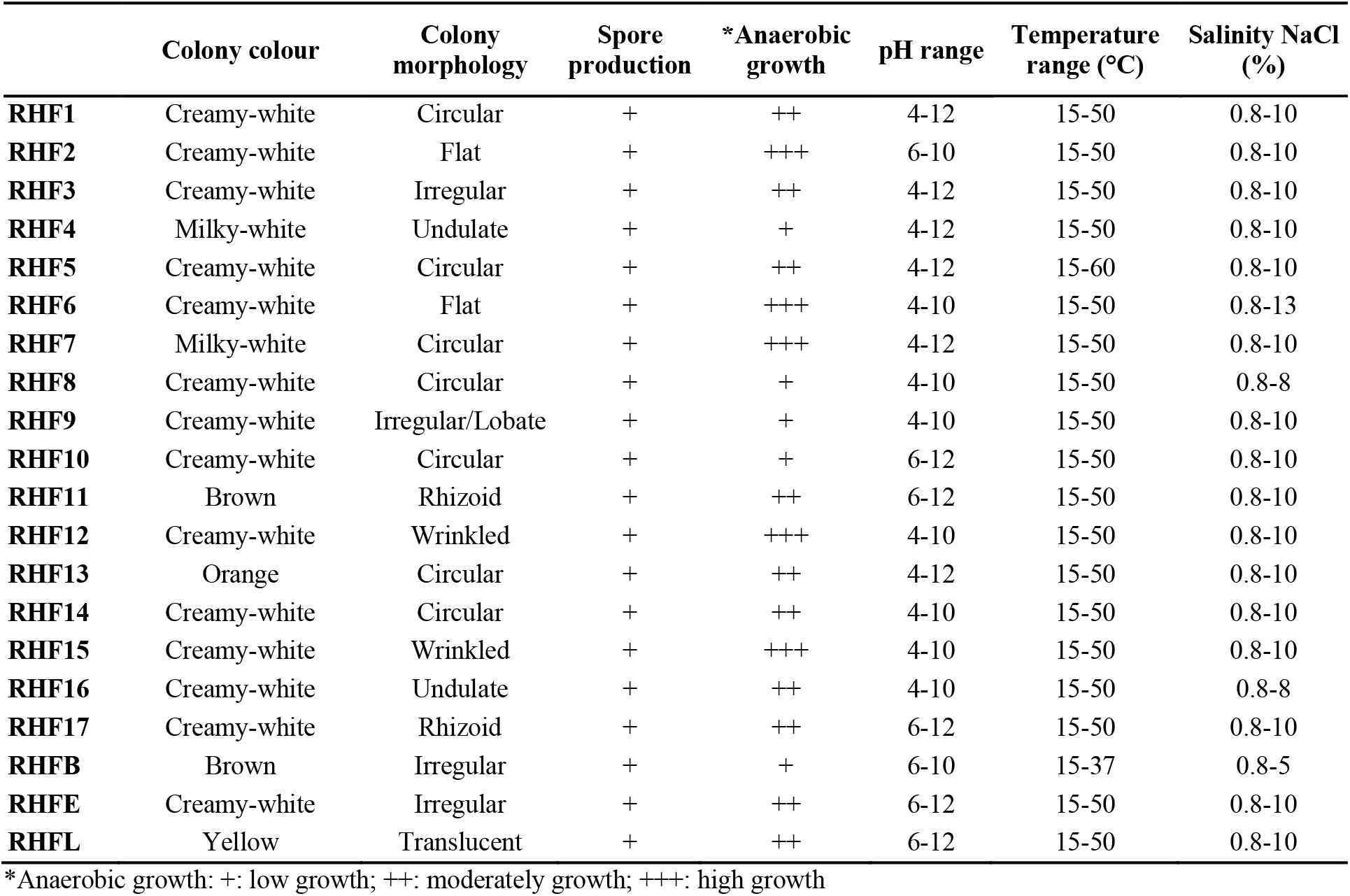
Preliminary characterization of spore-forming bacteria isolated from salt-pans.

**Table S3.** Pairwise average nucleotide identities (ANI) between the isolated strains and the closest relatives identified in the polyphasic analysis.

|  | **RHFB** | **RHF2** | **RHF6** | **RHF12** | **RHF15** | **RHFS10** | **RHFS18** |
| --- | --- | --- | --- | --- | --- | --- | --- |
| *B. subtilis* 168 | 68.21 | **99.96** | 77.28 | 87.34 | 98.79 | 91.84 | 76.88 |
| *B. gibsonii* | 68.47 | 98.83 | 77.09 | 87.3 | **99.6** | 91.76 | 77.01 |
| *B. amyloliquefaciens* | 67.94 | 77.1 | **99.26** | 77.17 | 76.83 | 77.13 | **98.36** |
| *B. velezensis* | 68.23 | 77.25 | 99.15 | 77.21 | 77.12 | 77.12 | 98.35 |
| *B. halotolerans* | 68.38 | 87.45 | 77.42 | **98.04** | 87.38 | 87.79 | 77.02 |
| *B. frigoritolerans* | **96.95** | 68.46 | 67.61 | 68.47 | 68.56 | 68.17 | 67.94 |
| *B. vallismortis* | 68.25 | 91.01 | 77.16 | 87.29 | 90.87 | **93.48** | 77.33 |

**Table S4.** Identified PGPR gene clusters and similarity (amino acid identity) to the closest organism (when available).

| Strain and gene cluster | Length (bp) | Type | Compound | Similarity (%) | Organism |
| --- | --- | --- | --- | --- | --- |
| **RHFB** |  |  |  |  |  |
| Cluster 1 | 24169 | NRPS | fengycin | 46 | *B. velezensis* FZB42 |
| Cluster 2 | 23535 | RiPP (LAP) | unknown |  |  |
| Cluster 3 | 20818 | terpene | unknown |  |  |
| Cluster 4 | 16393 | RiPP | paeninodin | 100 | *Paenibacillus dendritiformis* C454 |
| Cluster 5 | 21895 | terpene | unknown |  |  |
| Cluster 6 | 15513 | siderophore | unknown |  |  |
| Cluster 7 | 41088 | T3PKS | unknown |  |  |
| Cluster 8 | 49726 | NRPS | koraminine | 87 | *B. sp. NK2003* |
| Cluster 9 | 43445 | NRPS | bacillibactin | 53 | *B. subtilis subsp. subtilis str. 168* |
| **RHF2** |  |  |  |  |  |
| Cluster 1 | 20518 | terpene | unknown |  |  |
| Cluster 2 | 114759 | Hybrid PKS/NRPS | bacillaene | 100 | *B. velezensis* FZB42 |
| Cluster 3 | 72650 | NRPS | fengycin | 100 | *B. velezensis* FZB42 |
| Cluster 4 | 41097 | terpene | unknown |  |  |
| Cluster 5 | 20746 | CDPS | unknown |  |  |
| Cluster 6 | 49741 | NRPS | bacillibactin | 100 | *B. subtilis subsp. subtilis str. 168* |
| Cluster 7 | 65391 | NRPS | surfactin | 82 | *B. velezensis* FZB42 |
| Cluster 8 | 41418 | Other | bacilysin | 100 | *B. velezensis* FZB42 |
| Cluster 9 | 21611 | RiPP (Thiopeptide) | subtilisin A | 100 | *B. subtilis subsp. spizizenii* ATCC 6633 |
| Cluster 10 | 22953 | RiPP (Head-to-tailcyclized peptide) | sporulation killing factor | 100 | *B. subtilis subsp. subtilis str. 168* |
| Cluster 11 | 11461 | RRE-containing | unknown |  |  |
| **RHF6** |  |  |  |  |  |
| Cluster 1 | 105763 | Hybrid PKS/NRPS | difficidin | 100 | *B. velezensis* FZB42 |
| Cluster 2 | 40094 | T3PKS | unknown |  |  |
| Cluster 3 | 119121 | NRPS | fengycin | 93 | *B. velezensis* FZB42 |
| Cluster 4 | 109377 | Hybrid PKS/NRPS | bacillaene | 100 | *B. velezensis* FZB42 |
| Cluster 5 | 88230 | PKS | macrolactin H | 100 | *B. velezensis* FZB42 |
| Cluster 6 | 20740 | terpene | unknown |  |  |
| Cluster 7 | 66315 | NRPS | unknown |  |  |
| Cluster 8 | 51793 | NRPS | bacillibactin | 100 | *B. subtilis subsp. subtilis str. 168* |
| Cluster 9 | 41418 | Other | bacilysin | 100 | *B. velezensis FZB42* |
| Cluster 10 | 65407 | NRPS | surfactin | 82 | *B. velezensis FZB42* |
| Cluster 11 | 41244 | Saccharide | butirosin A/ butirosin B | 7 | *B. circulans* |
| Cluster 12 | 40884 | Phosphonate | unknown |  |  |
| **RHF12** |  |  |  |  |  |
| Cluster 1 | 60718 | NRPS | surfactin | 82 | *B. velezensis FZB42* |
| Cluster 2 | 41097 | T3PKS | unknown |  |  |
| Cluster 3 | 21612 | RiPP (Thiopeptide) | subtilisin A | 100 | *B. subtilis subsp. spizizenii ATCC 6633* |
| Cluster 4 | 41418 | Other | bacilysin | 100 | *B. velezensis FZB42* |
| Cluster 5 | 95323 | Hybrid PKS/NRPS | bacillaene | 92 | *B. velezensis FZB42* |
| Cluster 6 | 49738 | NRPS | bacillibactin | 100 | *B. subtilis subsp. subtilis str. 168* |
| Cluster 7 | 62913 | NRPS | fengycin | 93 | *B. velezensis FZB42* |
| Cluster 8 | 14447 | terpene | unknown |  |  |
| Cluster 9 | 11709 | terpene | unknown |  |  |
| Cluster 10 | 5195 | lanthipeptide-classIII | unknown |  |  |
| Cluster 11 | 2854 | RiPP -like | unknown |  |  |
| **RHF15** |  |  |  |  |  |
| Cluster 1 | 40888 | T3PKS | unknown |  |  |
| Cluster 2 | 20849 | Terpene | unknown |  |  |
| Cluster 3 | 82212 | NRPS | fengycin | 100 | *B. velezensis FZB42* |
| Cluster 4 | 114771 | Hybrid PKS/NRPS | bacillaene | 100 | *B. velezensis FZB42* |
| Cluster 5 | 20803 | Terpene | unknown |  |  |
| Cluster 6 | 41418 | Other | bacilysin | 100 | *B. velezensis FZB42* |
| Cluster 7 | 21611 | RiPP (Thiopeptide) | subtilisin A | 100 | *B. subtilis subsp. spizizenii ATCC 6633* |
| Cluster 8 | 20746 | CDPS | unknown |  |  |
| Cluster 9 | 49741 | NRPS | bacillibactin | 100 | *B. subtilis subsp. subtilis str. 168* |
| Cluster 10 | 65391 | NRPS (Lipopeptide) | surfactin | 82 | *B. velezensis FZB42* |
| Cluster 11 | 24457 | RiPP (Lanthipeptide) | subtilomycin | 100 | *B. subtilis* |
| **RHFS10** |  |  |  |  |  |
| Cluster 1 | 41097 | T3PKS | unknown |  |  |
| Cluster 2 | 21898 | Terpene | unknown |  |  |
| Cluster 3 | 128639 | NRPS | fengycin | 100 | *B. velezensis* FZB42 |
| Cluster 4 | 114812 | Hybrid PKS/NRPS | bacillaene | 100 | *B. velezensis* FZB42 |
| Cluster 5 | 41418 | Other | bacilysin | 100 | *B. velezensis* FZB42 |
| Cluster 6 | 21613 | RiPP (Thiopeptide) | subtilisin A | 100 | *B. subtilis* subsp. *spizizenii* ATCC 6633 |
| Cluster 7 | 20746 | CDPS | unknown |  |  |
| Cluster 8 | 49742 | NRPS | bacillibactin | 100 | *B. subtilis* subsp. *subtilis* str. 168 |
| Cluster 9 | 81352 | Hybrid PKS/NRPS | zwitermicin A | 18 | *B. cereus* |
| Cluster 10 | 20788 | Terpene | unknown |  |  |
| Cluster 11 | 65394 | PKS | macrolactin H | 90 | *B. velezensis* FZB42 |
| Cluster 12 | 22952 | RiPP (Head-to-tailcyclized peptide) | sporulation killing factor | 100 | *B. subtilis* subsp. *subtilis* str. 168 |
| **RHFS18** |  |  |  |  |  |
| Cluster 1 | 105749 | Hybrid PKS/NRPS | Difficidin | 100 | *B. velezensis* FZB42 |
| Cluster 2 | 40665 | T3PKS | unknown |  |  |
| Cluster 3 | 20138 | Terpene | unknown |  |  |
| Cluster 4 | 120565 | NRPS | fengycin | 93 | *B. velezensis* FZB42 |
| Cluster 5 | 109609 | Hybrid PKS/NRPS | bacillaene | 100 | *B. velezensis* FZB42 |
| Cluster 6 | 88235 | PKS | macrolactin H | 100 | *B. velezensis* FZB42 |
| Cluster 7 | 20740 | Terpene | unknown |  |  |
| Cluster 8 | 41244 | Saccharide | butirosin A/ butirosin B | 7 | *B. circulans* |
| Cluster 9 | 65410 | NRPS (Lipopeptide) | surfactin | 82 | *B. velezensis* FZB42 |
| Cluster 10 | 29736 | LAP, thiopeptide | unknown |  |  |
| Cluster 11 | 8 | Other | bacilysin | 100 | *B. velezensis* FZB42 |
| Cluster 12 | 51794 | NRPS | bacillibactin | 100 | *B. subtilis* subsp. *subtilis* str. 168 |

